# Divergence of gene regulatory network linkages during specification of ectoderm and mesoderm in early development of sea urchins

**DOI:** 10.1101/044149

**Authors:** Eric M. Erkenbrack, Eric H. Davidson

## Abstract

Developmental gene regulatory networks (GRNs) are assemblages of gene regulatory interactions that direct ontogeny of animal body plans. Studies of GRNs operating in early development of euechinoid sea urchins has revealed that little appreciable change has occurred since their divergence approximately 90 million years ago (mya). These observations suggest that strong conservation of GRN architecture has been maintained in early development of the sea urchin lineage. To test whether this is true for all sea urchins, comparative analyses of echinoid taxa that diverged deeper in geological time must be conducted. Recent studies highlighted extensive divergence of skeletogenic mesoderm specification in the sister clade of euechinoids, the cidaroids, suggesting that comparative analyses of cidaroid GRN architecture may confer a greater understanding of the evolutionary dynamics of developmental GRNs. Here, we report spatiotemporal patterning of 55 regulatory genes and perturbation analyses of key regulatory genes involved in euechinoid oral-aboral patterning of non-skeletogenic mesodermal and ectodermal domains in early development of the cidaroid *Eucidaris tribuloides*. Our results indicate that developmental GRNs directing mesodermal and ectodermal specification have undergone marked alterations since the divergence of cidaroids and euechinoids. Notably, statistical and clustering analyses of echinoid temporal gene expression datasets indicate that regulation of mesodermal genes has diverged more markedly than regulation of ectodermal genes. Although research on indirect-developing euechinoid sea urchins suggests strong conservation of GRN circuitry during early embryogenesis, this study indicates that since the divergence of cidaroids and euechinoids developmental GRNs have undergone significant divergence.

## Significance Statement

Sea urchins consist of two subclasses, cidaroid and euechinoids. Research on gene regulatory networks (GRNs) in early development of three euechinoids indicates that little appreciable change has occurred to their linkages since they diverged approximately 90 million years ago (mya). We asked if this conservation extends to the all echinoids. Here we demonstrate extensive divergence of GRN architecture in early embryonic specification of the oral-aboral axis in echinoids. We systematically analyzed in a cidaroid spatiotemporal expression and function of regulatory genes segregating euechinoid ectoderm and mesoderm. While we found evidence for diverged regulation of both mesodermal and ectodermal genes, our comparative analyses indicated that, since these two clades diverged 268.8 million years ago, mesodermal GRNs have undergone significantly more alteration than ectodermal GRNs.

\body

## Introduction

Integral to early development of a bilaterian is the development of the three embryonic tissue-layer domains: endoderm, ectoderm, and mesoderm. Asymmetrically distributed RNA and proteins in the egg provide the initial inputs into this process and thereby determine the spatial coordinates of domain formation (1, 2). In the context of these maternal factors, zygotic transcription of distinct sets of regulatory genes occurs in specific regions of the embryo, initiating the genomically encoded regulatory program and its output of regulatory genes. In this way the embryo becomes populated by distinct sets of transcription factors and cell signaling transcripts called regulatory states (3). The spatial readout of developmental gene regulatory networks (GRNs), regulatory states provide each cell with its molecularly distinct and functional identity (4, 5).

Sea urchins (class Echinodea) have long served as model systems to study mechanisms of early development and, more recently, to study fundamental aspects of developmental GRNs. Diversification of cell lineages and embryonic domains in sea urchin embryos depends on the cleavage positions of their blastomeres (6, 7), facilitating interpretation of mechanisms of spatial change. Sea urchin lineages have undergone multiple changes in life history strategies, providing a convenient framework, with experimental replicates, to investigate evolution and mechanisms of developmental programs (8). Importantly, a well-represented fossil record affords dating of evolutionary events (9) and has established that the sister subclasses of sea urchins—cidaroids and euechinoids—diverged from one another at least 268.8 million years ago (mya) (10). Differences in the timing of developmental events in embryogenesis of cidaroids and euechinoids have long been a topic of interest, but only recently have become the subject of molecular research (11–18).

Research on early development of the euechinoid purple sea urchin *Strongylocentrotus purpuratus* has brought into high resolution the players and molecular logic directing developmental GRNs that specify its varied embryonic domains and subdomains (19–30). Additionally, abundant comparative evidence exists for other euechinoid taxa, including *Lytechinus variegatus* (31–37) and *Paracentrotus lividus* (38–42). Data from these three indirect-developing euechinoids indicate that, although these taxa diverged from one another approximately 90 mya (9, 43), very little appreciable change to developmental GRNs has been observed in their early development (44–46). While there is evidence of a few heterochronic alterations to these GRNs, e.g. a heterochronic shift in *snail* expression in *L. variegatus* versus *S. purpuratus* (22), numerous studies have made clear the striking conservation of GRN linkages and spatiotemporal gene expression in these lineages. Recently, studies of early development of the distantly-related, indirect-developing cidaroid sea urchin *Eucidaris tribuloides* revealed that skeletogenic mesoderm specification in this clade is markedly different from that observed in euechinoids (17, 18, 47). These observations suggest that comparative analyses of GRN circuitry in early development of cidaroids and euechinoids have the potential to illuminate the tempo and mode of evolution of developmental GRNs.

Here, we coupled comparative spatiotemporal gene expression analyses with experimental manipulations of regulatory genes specifying euechinoid non-skeletogenic mesodermal (NSM) and ectodermal domains in the cidaroid *E. tribuloides* to reveal how these GRNs have changed since the divergence of these two clades 268.8 mya. This study focused on oral-aboral (O-A; also called dorsal-ventral) patterning, which has consequences for both ectoderm and mesoderm. O-A axis specification is a well-documented process in euechinoids (46) and is a highly conserved developmental mechanism in deuterostomes (48, 49). We present evidence that deployment and regulatory interactions of regulatory genes specifying sea urchin ectoderm and mesoderm have diverged substantially in indirect-developing echinoids. Importantly, our comparative data and analyses suggest that regulatory interactions occurring in ectodermal GRNs have undergone less divergence than those occurring in mesodermal domains. Our conclusions are supported by comparative spatiotemporal data, statistical analyses of timecourse gene expression data in three taxa of echinoids and perturbation analyses. Taken together regulatory genes expressed in ectodermal domains exhibited stronger signals of conservation relative to those expressed in mesodermal domains. These results suggest that embryonic domains and cell types in early development of sea urchins have evolved at different rates since the divergence of the two sister sub-classes of modern sea urchins. These alterations to GRN architecture have occured throughout the network and frequently since their divergence. Additionally, these results offer an in principle explanation for observations of rapid change to nearly all components of developmental process in the development of direct-developing, non-feeding sea urchins (50–53).

## Results

### Ectodermal O-A regulatory states in the cidaroid *E. tribuloides* suggest striking conservation of regulatory gene deployment in echinoids

In euechinoids, ectoderm is segregated into a diverse set of regulatory states defined by the future location of the stomodeum (46, 54, 55). A critical factor in establishing oral-aboral (47) polarity in sea urchins is the signaling ligand *nodal* (35, 38). Nodal ligand activates the SMAD signaling pathway, which is directly upstream of *nodal* (itself), *not, lefty* and *chordin* in oral ectoderm (OE) (40, 56). In *E. tribuloides,* zygotic transcription of *nodal, not* and *lefty* begins by early blastula stage (Fig. 1*D*, 1*J*, and Fig. S1*D*). In contrast, transcriptional activation of *chordin* is delayed by at least 5 hours from this initial cohort, indicative of an intermediate regulator between *nodal* and *chordin* in *E. tribuloides* (Fig. 1D and 1E). From 17 hpf to 40 hpf, spatial expression of *nodal* is observed in a well-defined region in OE that expands slightly as gastrulation proceeds (Fig. 1D2–1D5 and Fig. S2). Unlike *nodal*, the spatial distribution of its targets is not solely restricted to a small field of cells in OE. *Lefty* (also known as Antivin), an antagonist of *nodal*, exhibits a broader pattern of expression that, by 50 hpf, expands into the oral side of the archenteron (Fig. S1D2-S1D5 and Fig. S2). Similarly, *chordin* transcripts are detected in OE throughout early *E. tribuloides* development (Fig. 1E1–1E5 and Fig. S2). The homeobox gene *not,* known to play a role directly downstream of *nodal* in euechinoid O-A ectodermal and mesodermal polarization (27, 57), was observed spatially in OE during gastrulation and later extends vegetally towards the perianal ectoderm and is observed in the archenteron (Fig. 1J1–1J5 and Fig. S2). In euechinoids, the *bmp2/4* ligand is a directly downstream of Nodal and is translocated across the embryo to the aboral side, where it stimulates via SMAD signaling in aboral ectoderm (AE) specification genes such as *tbx2/3* (38, 58, 59). In *E. tribuloides*, both *bmp2/4* and *tbx2/3* are upregulated with the *nodal-not-lefty* cohort (Fig. S1G and Fig. 1I1–1I5). *Tbx2/3* exhibits spatial expression from late blastula stage onwards that is complementary to OE genes in the lateral AE (Fig. 1I2–1I5). By mid-gastrula stage, *tbx2/3* is also expressed in the archenteron and much later, by 70 hpf, is expressed in the bilateral clusters of cells synthesizing the larval skeleton (Fig. S2), which is similar to the spatial expression in two euechinoids with notably interesting heterochronic differences (60, 61). Lastly, the Forkhead family transcription factor *foxq2* is sequentially restricted to and specifically expressed in embryonic anterior neural ectoderm (ANE) territory in deuterostomes (62). In euechinoids, *foxq2* restriction to ANE is a crucial component of O-A axis specification, setting the anterior boundary of OE by restricting expression of *nodal* (23, 55). In *E*. *tribuloides, foxq2* exhibited an expression pattern consistent with observations in euechinoids and other deuterostomes, suggesting conserved roles for this gene in ANE and O-A specification (Fig. 1B and Fig. S2).

**Fig 1.**
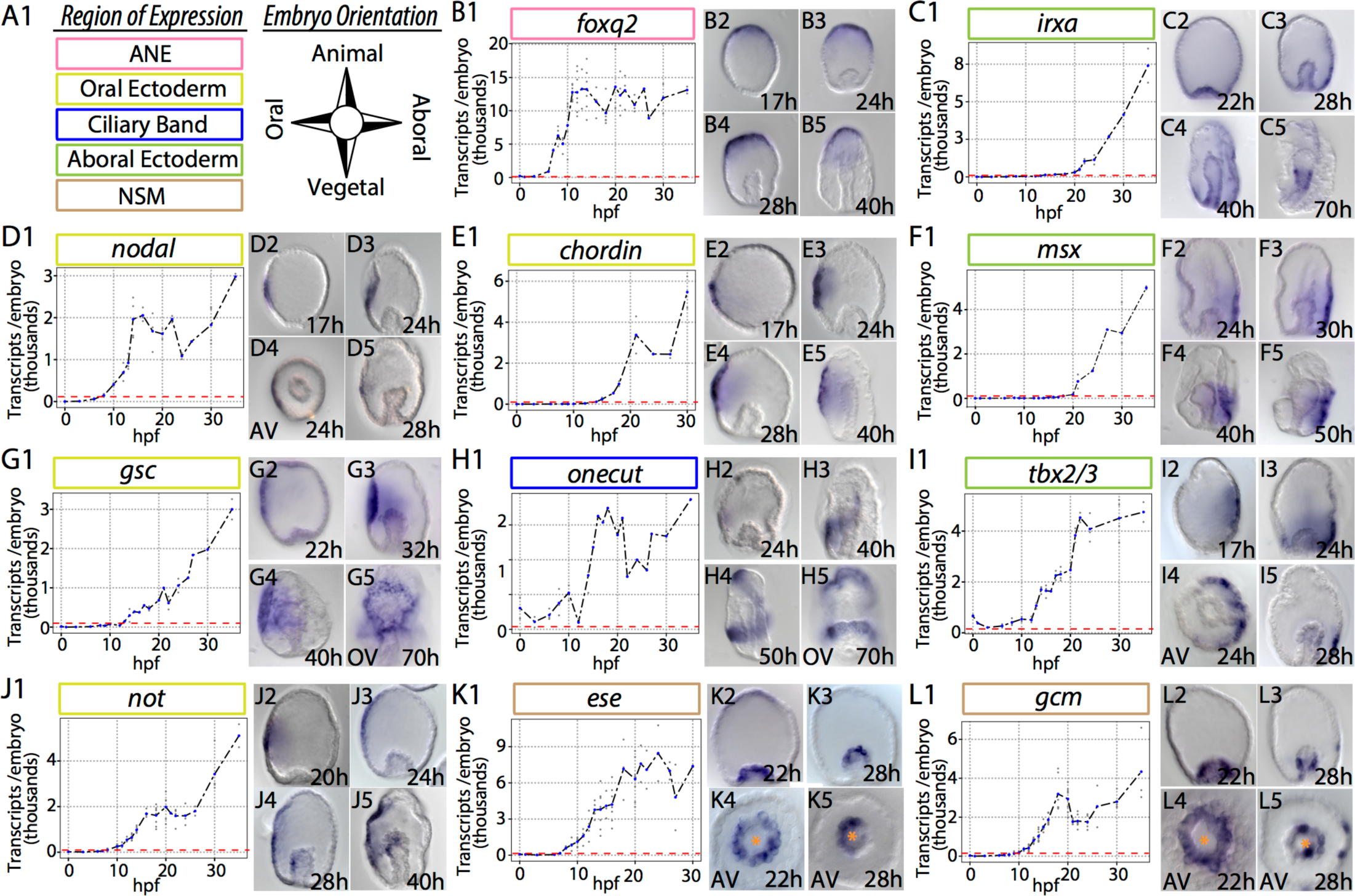
Spatiotemporal dynamics of eleven euechinoid ectodermal and mesodermal regulatory genes in the cidaroid *E. tribuloides*. Visualization of mRNA transcripts revealed by whole mount in situ hybridization, and estimates of absolute mRNA transcript abundance determined by qPCR during first 35 hours post fertilization (hpf). Individual data points are light grey. Blue data points represent the mean at that particular timepoint. Red dashed lines indicate the onset of zygotic transcription. Orange asterisks denote position of archenteron. (A1) Key to embryonic domains of expression and orientation of embryo micrographs. The embryonic domain of expression of each gene is indicated by colored boxes: ANE (anterior neural ectoderm), pink; oral ectoderm (OE), yellow; ciliary band (CB), blue; aboral ectoderm (AE), green; NSM (non-skeletogenic mesoderm), brown. (B1-B5) Spatiotemporal expression of *foxq2. Foxq2* spatial distribution is detected prior to hatching and is spatially restricted to ANE by 17 hpf. (C1-C5) Spatiotemporal expression of *irxa*. At 28 hpf, *irxa* is detected in AE and extends from the vegetal endodermal domains to ANE. By 40 hpf *irxa* is seen extending anteriorly at the boundary of AE and OE. (D1-D5) Spatiotemporal expression of *nodal*. *Nodal* spatial distribution is restricted to a few cells in OE up to early-mid gastrula stage. (E1-E5) Spatiotemporal expression of *chordin*. First observed in a few cells in OE at 17 hpf, *chordin* subsequently expands to extend from the perianal ectoderm to ANE. (F1-F5) Spatiotemporal dynamics of *msx*. Detected exclusively in AE, *msx* extends from the blastopore to ANE. (G1-G5) Spatiotemporal expression of *gsc*. By 22 hpf, *gsc* is detected in a broad region surrounding OE and later is observed around the stomodeum. (H1-H5) Spatiotemporal dynamics of *onecut*. By 40 hpf *onecut* is detected in the future post oral ciliary band and is initiated in a band moving from the posterior to the anterior. (I1-I5) Spatiotemporal expression of *tbx2/3*. Spatial distribution of *tbx2/3* at 17 hpf is detected broadly in aboral ectoderm (AE) and later extends from the perianal ectoderm to lateral AE, but not past the embryonic equator. (J1-J5) Spatiotemporal expression of *not. Not* spatial distribution is first detected in a similar field of cells as *nodal* and *lefty;* however, the domain of *not* subsequently expands by 28 hpf where it is seen in the oral side of the archenteron, where non-skeletogenic mesoderm (NSM) and endoderm are being segregated. By 40 hpf, the spatial domain of *not* extends from anterior neural ectoderm to the perianal ectoderm, is clearly seen in NSM, and was not detected in endodermal lineages. (K1-K5) Spatiotemporal expression of *ese. Ese* is detected primarily in non-skeletogenic mesoderm (NSM) but also in anterior neural ectoderm. By 28 hpf, *ese* is observed at the tip of the archenteron and is asymmetrically polarized in NSM. (L1-L5) Spatiotemporal expression of *gcm*. As gastrulation begins *gcm* is expressed broadly in NSM; however by 28 hpf it exhibits stark polarity in a field of cells that resides basal to the tip of the archenteron. *Gcm* is also observed in a few ectodermal cells at the time of primary mesenchymal ingression.

### Conserved spatiotemporal deployment of ciliary band regulatory genes

Free-feeding, indirect-developing sea urchins possess a single neurogenic ciliated band (CB) early in development that circumnavigates the larval oral face and facilitates feeding and locomotion (63). This structure has undergone frequent modification in the lineages leading to modern sea urchins, viz. in planktotrophic larvae (64). In euechinoids *goosecoid (gsc), onecut* and *irxa* contribute to the geometric patterning of CB formation (28, 40, 65). In euechinoids, *gsc* is expressed in OE and is directly downstream of *nodal* signaling on the oral side of the embryo (40); *onecut* (also known as *hnf6*) is a ubiquitous, maternally deposited factor that is restricted to a region spanning the boundaries of OE and AE, at which lies progenitor CB territory; and *irxa* is expressed exclusively in AE downstream of *tbx2/3* (40, 66). In the cidaroid *E. tribuloides*, *gsc* is activated with the OE cohort and is also spatially restricted to OE (Fig. 1G1 and Fig. S3). As in euechinoids, *onecut* is maternally deposited in *E. tribuloides* (Fig. 1H1). Although its early spatial expression was not obtained, staining was observed in a restricted band of cells encircling the OE by mid-gastrula stage. The spatial dynamics of *onecut* in *E. tribuloides* is notable insofar as its spatial distribution in progenitor CB begins in the progenitor field of post oral CB and subsequently extends in a narrow band of 4-8 cell diameters towards progenitor pre oral CB (Fig. 1H2–1H5 and Fig. S3). This observation starkly contrasts that in euechinoids, in which *onecut* is observed ubiquitously expressed early and later delimited as a whole to the CB territory by transcriptional repressors in the OE and AE (28, 67). *Irxa* initiates zygotic expression at mid-blastula stage (∼14 hpf) in *E. tribuloides*, and by 28 hpf is observed broadly in AE (Fig. 1C1–1C5). Unlike in euechinoids, *irxa* is broadly distributed in *E. tribuloides* AE—much more so than *tbx2/3*—indicating that it is likely broadly activated in the ectoderm and repressed in OE and ANE. These spatiotemporal observations of key CB regulatory genes suggest that CB patterning mechanisms are likely to be conserved in echinoids.

### Non-skeletogenic mesodermal patterning and regulatory interactions have diverged widely in echinoids

Non-skeletogenic mesoderm (NSM) in euechinoids arises at the vegetal plate from early cleavage endomesodermal precursors and gives rise to four different cell types: blastocoelar cells, pigment cells, circumesophageal cells and coelomic pouch cells (68). Experimental observations indicate that euechinoids completely rely on presentation of Delta ligand in the adjacent SM to upregulate NSM regulatory factors in veg2 endomesodermal cells (32, 69, 70). After Delta/Notch signaling upregulates NSM regulatory genes in SM-adjacent cells, NSM is partitioned into aboral NSM and oral NSM as a result of Nodal/SMAD signaling upregulating transcription factor *not* on the oral side of the embryo (27, 71). The first evidence of NSM polarity in *E. tribuloides*, is seen in the spatial distribution of *ese* and *gcm* transcripts shortly after gastrulation has commenced (Fig. 1K2–1K5 and 1L2-1L5). In euechinoids, *ese* is expressed in the oral NSM, and *gcm* is expressed in the aboral NSM. In *E. tribuloides ese* is first in ANE and progenitor NSM just prior to the onset of gastrulation, shortly after invagination of the archenteron it is subsequently restricted to one side in NSM (Fig. 1K2–1K5). Similarly, *gcm* is expressed transiently in oral and aboral NSM, and by 28 hpf is restricted to a cluster of cells just below the tip of the archenteron (Fig. 1L2–1L5). Later this expression is seen solely on one side of the archenteron as *gcm-* positive cells ingress rapidly into the blastocoel at 36 hpf (Fig. S4). Double fluorescent whole-mount *in situ* hybridization (WMISH) revealed that *ese* and *gcm* in this cidaroid are restricted to opposite sides of the archenteron (Fig S4G1 and S4G2), providing the first evidence of conserved polarity in NSM segregation in echinoids.

To further clarify NSM regulatory states in *E. tribuloides* and their similarity to those in euechinoids, we conducted additional single and double WMISH on NSM regulatory genes *ets1/2, gatac, gatae, prox, scl* and *tbrain*. Directly downstream of *gcm* in euechinoids is *gatae* (27). In *S*. *purpuratus*, *gatae* is observed in endomesoderm by blastula stage (72). In *E. tribuloides* NSM, *gatae* is expressed throughout the endomesoderm at the time of SM ingression (∼28 hpf) and later is observed restricted to one side near the tip of the archenteron, as well as in the second wave of ingressing mesenchyme (Fig. S1C2-S1C5 and Fig. S4). *Gatac (gata1/2/3), prox* and *scl*, all of which are oral NSM genes in euechinoids (27), come off the baseline at similar times in *E*. *tribuloides* and are detectable in a few cells at the base of the vegetal pole by 18 hpf by WMISH (Fig. S4B1, S4E1, and S1F). Of these three genes, *scl* was the first to show O-A NSM polarity followed by *gatac* (Fig. S1B5, S1E5, and S1F5). Surprisingly, by 36 hpf *prox* did not exhibit spatial distribution of transcripts that unambiguously indicated O-A polarity (Fig. S1E2-S1E5 and Fig. S4E6), suggesting that either *prox* is a general mesodermal regulatory factor in *E. tribuloides* or it is spatially restricted later in gastrular development.

These data on spatial dynamics of NSM regulatory factors suggest that there exist numerous regulatory states in the anterior archenteron. To provide some clarity, double fluorescent WMISH (dfWMISH) indicated this was indeed the case. Previous observations suggested that *E*. *tribuloides* mesodermal domains broadly express *ets1/2* and *tbrain* (17), and dfWMISH confirmed this result (Fig. S4IS4I). Within this broad *ets1/2-tbrain* domain, three regulatory states are identified (Fig. S4G-S4I): (1) orally localized *ets1/2, tbrain* and *ese*; (2) aborally localized *ets1/2, tbrain,* and *gcm*; and (3) an anteriorly localized micromere-descendant regulatory state at the tip of the archenteron of *ets1/2, tbrain, ese* and *alx1*. The only similarity between NSM regulatory states of *E. tribuloides* and that of euechinoids is *ese* is restricted to oral NSM and *gcm* to aboral NSM (27). The NSM regulatory states described here make it clear that since the divergence of cidaroids and euechinoids there have been numerous changes to spatial and temporal gene expression during the early patterning of echinoid NSM cell types.

### Perturbation of Nodal signaling suggests highly conserved regulatory interactions in patterning echinoid O-A ectoderm and mesoderm

Next we sought to experimentally perturb O-A specification in *E. tribuloides* to determine the extent to which this developmental program is conserved in echinoids. In euechinoids, perturbation of animal-vegetal (A-V) axis polarity by disruption of nuclearization of β-catenin indirectly disrupts O-A axis specification (31, 33). One mechanism underlying the crosstalk of these two deuterostome specification events was found to be restriction of *foxq2* to ANE, as its presence in OE blocked *nodal* transcription (23). In *E. tribuloides,* disrupting nuclearization of β-catenin at the vegetal pole by overexpressing dn-Cadherin RNA led to upregulation of *foxq2* and strong downregulation of *nodal* and its euechinoid downstream components *bmp2/4, not,* and *tbx2/3* (Fig. S5A). This result suggests that the molecular crosstalk between β-catenin/TCF, *foxq2* and *nodal* are conserved in echinoids.

Next, we aimed to determine the spatiotemporal effects of perturbation of O-A specification by culturing *E. tribuloides* embryos in the presence of SB43152, a small molecule antagonist of the TGF-β (Nodal) receptor Alk4/5/7 (73). At four days post fertilization, these embryos exhibited strong aboralization, archenterons that failed to make contact with OE, and supernumerary skeletal elements (Fig. S5B). Quantitative PCR (qPCR) analysis at four different timepoints in *E. tribuloides* development showed strong downregulation of OE regulatory factors *chordin, gsc, lefty, nodal* and *not* (Fig. 2A). This result was confirmed spatially by WMISH for mRNA transcripts of *chordin, nodal* and *not* (Fig. 2B and S5C). Another critical OE regulatory gene is the secreted TGF-β ligand *bmp2/4*. This gene was clearly not affected to the same degree as the aforementioned cohort of OE factors (Fig. 2A). This result is strikingly different from the strong downregulation of *bmp2/4* observed in the euechinoid *P. lividus* when it was cultured in the presence of SB431542 or when injected with Nodal morpholino (MASO) (38, 40), indicating that regulation of *bmp2/4* may be under alternative control in *E. tribuloides*.

**Fig 2.**
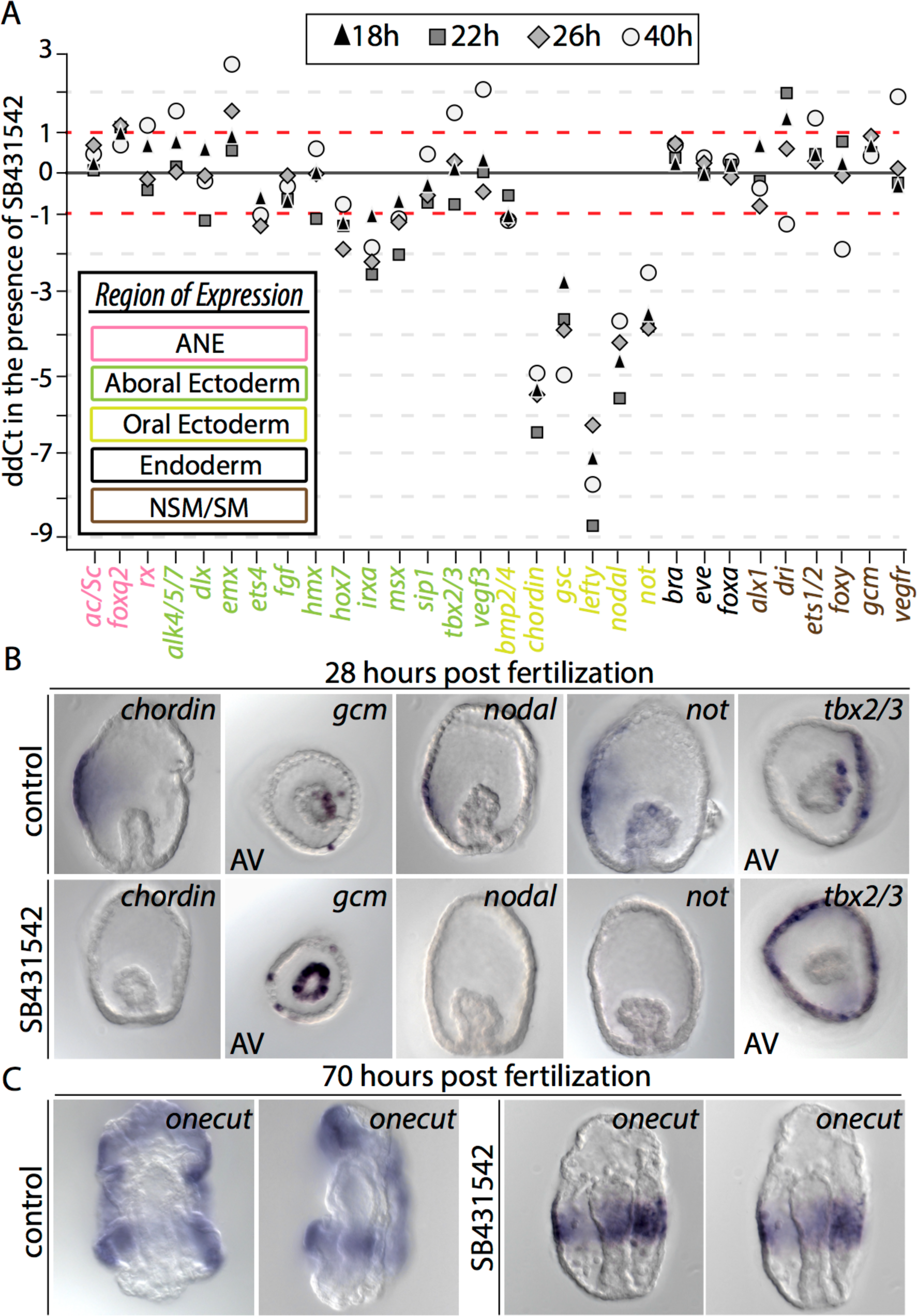
Perturbation of oral-aboral axis formation in *E. tribuloides* reveals alterations to regulatory factor deployment in echinoids. (A1) Quantitative effect of SB431542 on expression of 30 *E. tribuloides* regulatory genes as revealed by qPCR. Fold change in mRNA transcripts (ddCt) is shown on the y-axis. Two timepoints from two independent replicates are shown. Regulatory factors are listed on the x-axis and font color designates their embryonic domain: pink, anterior neural ectoderm; green, aboral ectoderm; yellow, oral ectoderm; black, endoderm; brown, mesoderm. (B) WMISH of select ectodermal and mesodermal regulatory genes in embryos cultured in the presence of SB431542. At 28 hpf expression of *chordin, nodal* and *not* are completely extinguished. Whereas *gcm* is regularly restricted to one side of the archenteron, in the presence of SB431542 it exhibits expression throughout the archenteron. In the ectoderm, expression *tbx2/3* expands from aboral ectoderm (AE) into oral ectoderm in the presence of the inhibitor. (C) The ciliary band marker *onecut* is normally observed in a band of cells between the boundaries of OE and AE. However, in the presence of SB431542, *onecut* is expressed in an equatorial band that is 6-10 cell diameters across.

On the aboral side, a striking difference is the effect of SB431542 on *tbx2/3*. In euechinoids, *tbx2/3* is downstream of *bmp2/4* ligand, which diffuses from OE to AE (58, 59, 74). Treatment of *P*. *lividus* embryos with SB431542 inhibitor completely and specifically extinguishes *tbx2/3* in AE while not interfering with its SM expression (40). In *E. tribuloides,* qPCR data suggest SB431542 inhibitor has no effect on *tbx2/3* regulation (Fig. 5C). However, WMISH of *tbx2/3* in the presence of the inhibitor revealed its domain of expression had expanded into an equatorial ring (Fig. 2B and S5C). Similarly, whereas in *E. tribuloides* qPCR data indicate strong downregulation of the AE regulatory factor *irxa* (Fig. 5C), its domain of expression expanded into OE in *P. lividus* embryos upon SB431542 treatment (40). These results suggest GRN architecture immediately downstream of the initial *nodal* and *bmp2/4* circuitry has diverged in echinoids and further indicate that regulatory interactions immediately upstream of *bmp2/4, irxa* and *tbx2/3* are regulated differently in *E. tribuloides*.

Between the OE and AE is the pro-neural ciliary band that expresses *onecut*. In euechinoids, perturbation of Nodal signaling results in an expansion of *onecut* throughout the ectoderm, leading to the suggestion that the default fate of ectoderm is a pro-neural CB territory (40). When this perturbation was carried out in *E. tribuloides,* a very different result was obtained in which we see *onecut* restricted to a single equatorial band of 6-10 cell diameters (Fig. 2C). A previous study showed that disruption of *E. tribuloides* endomesoderm formation by treatment with zinc results in embryos exhibiting a ring of highly concentrated proneural synaptotagmin-B positive cells at the equator of the embryo (75). Together, these observations indicate that, in the absence of Nodal signaling, there are anteriorly positioned repressors in *E. tribuloides* restricting CB fate to the equator of the embryo and that CB restriction from the ANE is functioning differently in cidaroids and euechinoids.

Lastly, in euechinoids studied thus far, polarity in NSM (47) lineages is downstream of Nodal signaling via the transcription factor Not (27, 71). In *E. tribuloides,* qPCR data did not indicate consistent differences in mRNA abundance for NSM regulatory genes (Fig. 2A). However, WMISH assays revealed that embryos treated with SB431542 failed to restrict *gcm* to the aboral side (Fig. 2B and S5C). This observation is consistent with the euechinoid GRN linkage immediately downstream of Nodal signaling via the transcription factor *not,* which represses aboral NSM in the oral-facing region of the archenteron (27). Indeed, in *E. tribuloides, not* is seen in the archenteron throughout gastrulation (Fig. 1J3–1J5). These observations are consistent with a conserved role for Nodal signaling via *not* in establishing polarity in NSM cell types in the archenteron of *E. tribuloides*.

### Regulatory interactions establishing ciliary band are conserved in echinoids

The spatial distributions of *gsc, onecut* and *irxa* are highly suggestive of a conserved regulatory apparatus that spatially restricts CB to the boundary of OE and AE (Fig. 1 and Fig. S3). Supporting this claim, microinjection of mutated or wild-type BACs encoding the *cis-* regulatory region of *S. purpuratus onecut* and a GFP reporter were examined in *E. tribuloides*. In *S*. *purpuratus, onecut* is restricted to the progenitor CB territory by *gsc* repression in OE and *irxa* repression in AE (Fig. 3A). Recently, Barsi and Davidson (2016) experimentally validated four cis-regulatory modules directing the activation of *onecut* by *soxb1* and repression of *onecut* by *gsc* in OE and *irxa* in AE (Fig. 3B). Remarkably, these BACs faithfully recapitulated the regulatory activity of these BACs in *E. tribuloides* (Fig. 3C–3H and Fig. S6). To test for this conservation, we assayed a series of endogenous and site-directed mutagenesis *onecut* BACs from *S. purpuratus* by microinjection (65). Remarkably, a BAC that has been shown to recapitulate the endogenous *S*. *purpuratus onecut* expression pattern faithfully expressed reporter GFP in the CB of *E. tribuloides* (Fig. 3H and S6A-S6B). Further, a BAC harboring mutated repressor sites for the oral repressor *gsc* repeatedly exhibited ectopic expression in OE of *E. tribuloides* (Fig. 3E). When interpreted with the spatial WMISH data of Figure 1, the early specification of CB regulatory factors suggests divergence of initial activation of *onecut* and conserved GRN circuitry guiding the restriction of *onecut* to CB territory by *irxa* and *gsc,* a striking example of a highly conserved developmental program after 268 million years of evolution.

**Fig 3.**
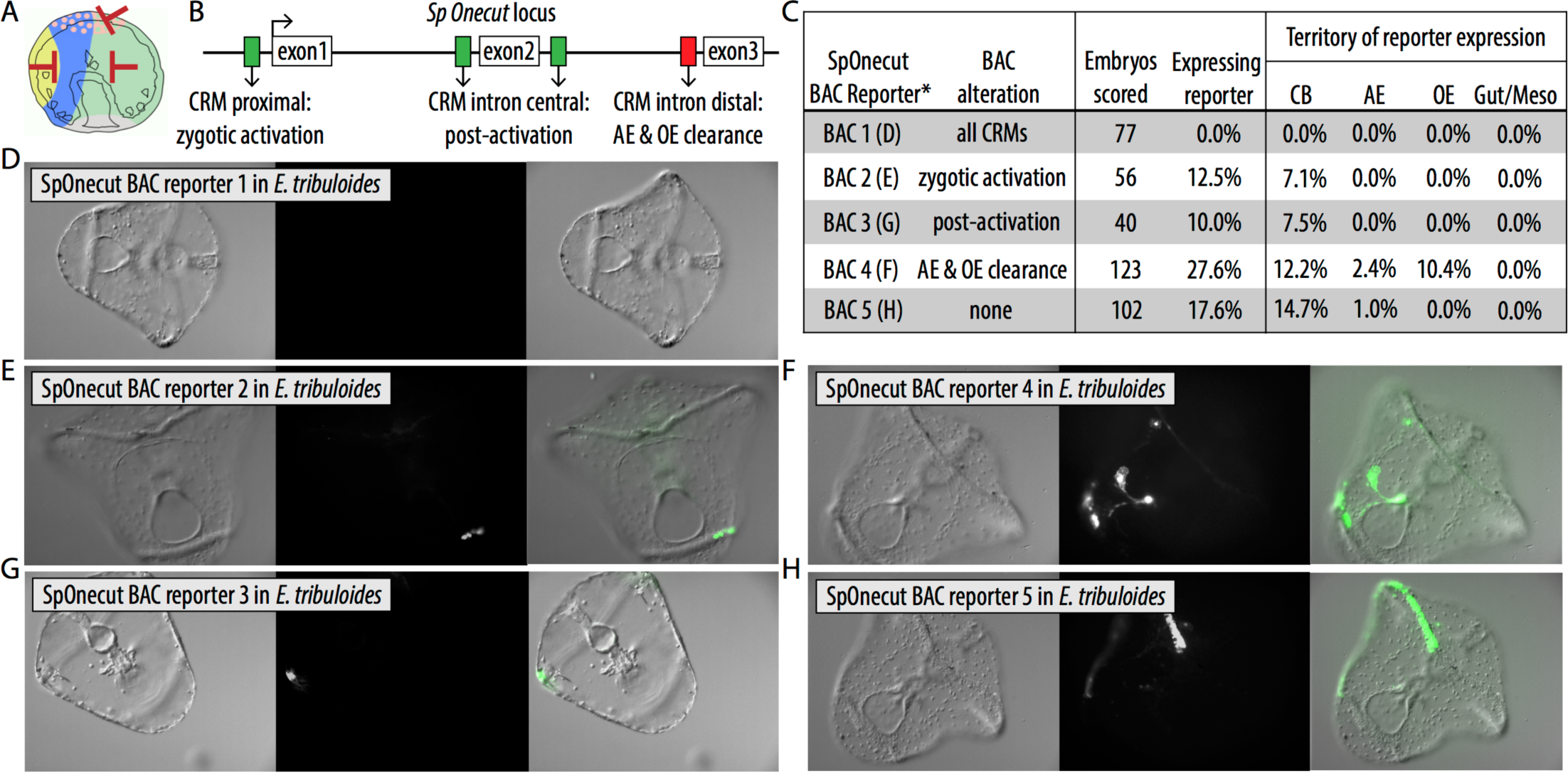
Expression dynamics in *E. tribuloides* of engineered reporter BACs harboring the regulatory locus of *S. purpuratus onecut* suggest conservation of GRN circuitry positioning ciliary band in echinoids. (A) Schematic of ciliary band (CB) restriction in the *S. purpuratus* embryo. Red bars indicate repressor genes restrict CB genes from expression in those domains. (B) Schematic showing the *S. purpuratus onecut* locus and the *cis-* regulatory modules (CRMs) responsible for the geometric positioning of *onecut*. (C) Table showing reporter analysis of spatial expression of five engineered *S. purpuratus onecut* BACs in *E. tribuloides*. The BACs were previously utilized to analyze the *cis*-regulatory dynamics of *onecut* spatial restriction in *S*. *purpuratus* (Barsi and Davidson, 2016). (D) BAC reporter 1, which harbors mutations to all known *cis-* regulatory modules (CRMs), showed a marked reduction of all reporter expression. (E) BAC reporter 2 harbors mutations to a CRM driving zygotic activation. (F) BAC Reporter 4, which harbors mutations to the aboral and oral ectodermal repression CRMs, increases the percent of embryos that show reporter expression in the oral ectoderm, as well as normal reporter expression in the ciliary band. (G) BAC reporter 3, which harbor mutated post-activation enhancer CRMs, showed reduced reporter expression both in terms of percent embryos exhibiting reporter activity and in the number of cells with reporter. (H) BAC Reporter 5, which harbors the unperturbed, wild-type locus of *S. purpuratus onecut,* faithfully exhibits reporter GFP in the ciliary band domain of *E*. *tribuloides*.

### Comparative analyses of temporal gene expression in echinoids suggests ectodermal and mesodermal GRNs have diverged at different rates

The spatiotemporal data presented thus far are highly suggestive that O-A axis specification, as well as gastrular CB formation, in *E. tribuloides* is consistent with similar processes in euechinoids and that NSM specification has ostensibly diverged. Next we sought a comparative gene expression approach to reveal the extent of divergence between echinoid clades. We compared absolute mRNA transcript abundance of 18 regulatory genes in three species and 55 *S. purpuratus* regulatory genes (76) in early euechinoid developmental GRNs for 14 early developmental timepoints in *E. tribuloides* (Table S5 and S6). We scaled timepoints from *E. tribuloides* to the euechinoid *P. lividus* since both develop at a similar temperature and since scaling of developmental stages between *P. lividus* and *S. purpuratus* had been previously established (Table S5) (77). This allowed pairwise analyses of Spearmann’s rank correlation coefficients, ρ, between the three species were conducted (Table S6). Plotting relative mRNA transcriptional dynamics for the three species were indicative of compelling correlation for ectodermal regulatory factors and supported the notion of poor correlation for regulatory factors driving mesoderm specification (Fig. 4 and S7). We next considered timecourse expression data of 55 orthologs of *E. tribuloides* and *S. purpuratus*. We binned the orthologs in ectoderm, endoderm and mesoderm based on their spatial distributions in *S. purpuratus* and compared them against the mean of all ρ values. Regulatory orthologs expressed in endoderm and ectoderm exhibited significantly higher ρ relative to the mean of all ρ values, suggesting strong conservation of transcriptional dynamics of these factors in echinoids (Fig. 5A). However, regulatory orthologs expressed in mesodermal cell lineages did not depart significantly from the mean ρ, suggesting transcriptional dynamics of mesodermal regulatory factors have changed markedly since the cidaroid-euechinoid divergence (Fig. 5A and 5B). Furthermore, we conducted a comparative clustering analysis to identify statistical differences in the timecourse data. By clustering *S*. *purpuratus* regulatory orthologs into six distinct clusters and forcing assignment of each *E*. *tribuloides* ortholog into all clusters, we acquired cluster membership scores and clustering similarities for each ortholog (Table S7, S8, and S9). Considering *S. purpuratus* orthologs alone revealed regulatory genes clustering together based on whether they are expressed maternal, early or late in development (Fig. S8A). When *E. tribuloides* orthologs were forced into these *S*. *purpuratus* clusters and the highest membership value for each ortholog was averaged with other orthologs expressed in each germ layer, *E. tribuloides* orthologs expressed in ectodermal domains exhibited higher cluster membership scores than orthologs expressed in mesodermal domains (Fig. 5C, p<0.05, Mann-Whitney U Test). Additionally, ectodermal orthologs jumped less frequently between clusters than mesodermal orthologs, suggesting timecourse data for ectodermal genes were more similar than those mesodermal genes (Fig. 5D and S8C). These analyses suggest that deployment of ectodermal regulatory genes is more similar between euechinoids and cidaroids than deployment of mesodermal regulatory genes. Further,

**Fig 4.**
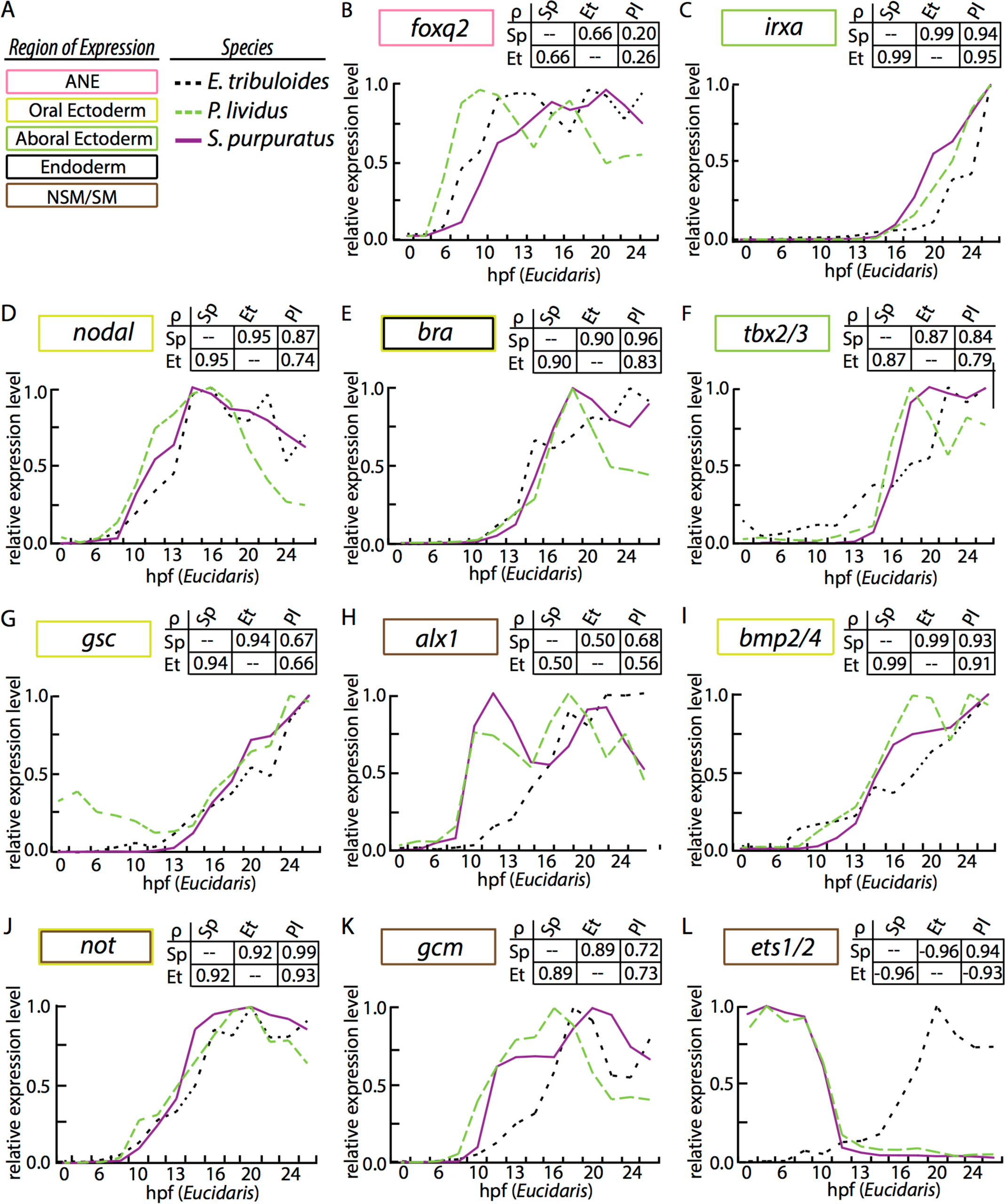
Comparative gene expression analysis of echinoid regulatory orthologs. *Strongylocentrotus purpuratus,* purple line *; Paracentrotus lividus,* green dashed line; *Eucidaris tribuloides,* black dashed line. Region of expression is indicated by a colored box. Transcripts per embryo for each gene were normalized to their maximal expression over the first 30 hours of development and are plotted against *E. tribuloides* development on the x-ordinate. Comparative developmental staging for each species is listed in Table S5. Each analysis is accompanied by a matrix of Spearman correlation coefficients (indicated as ρ).

**Fig 5.**
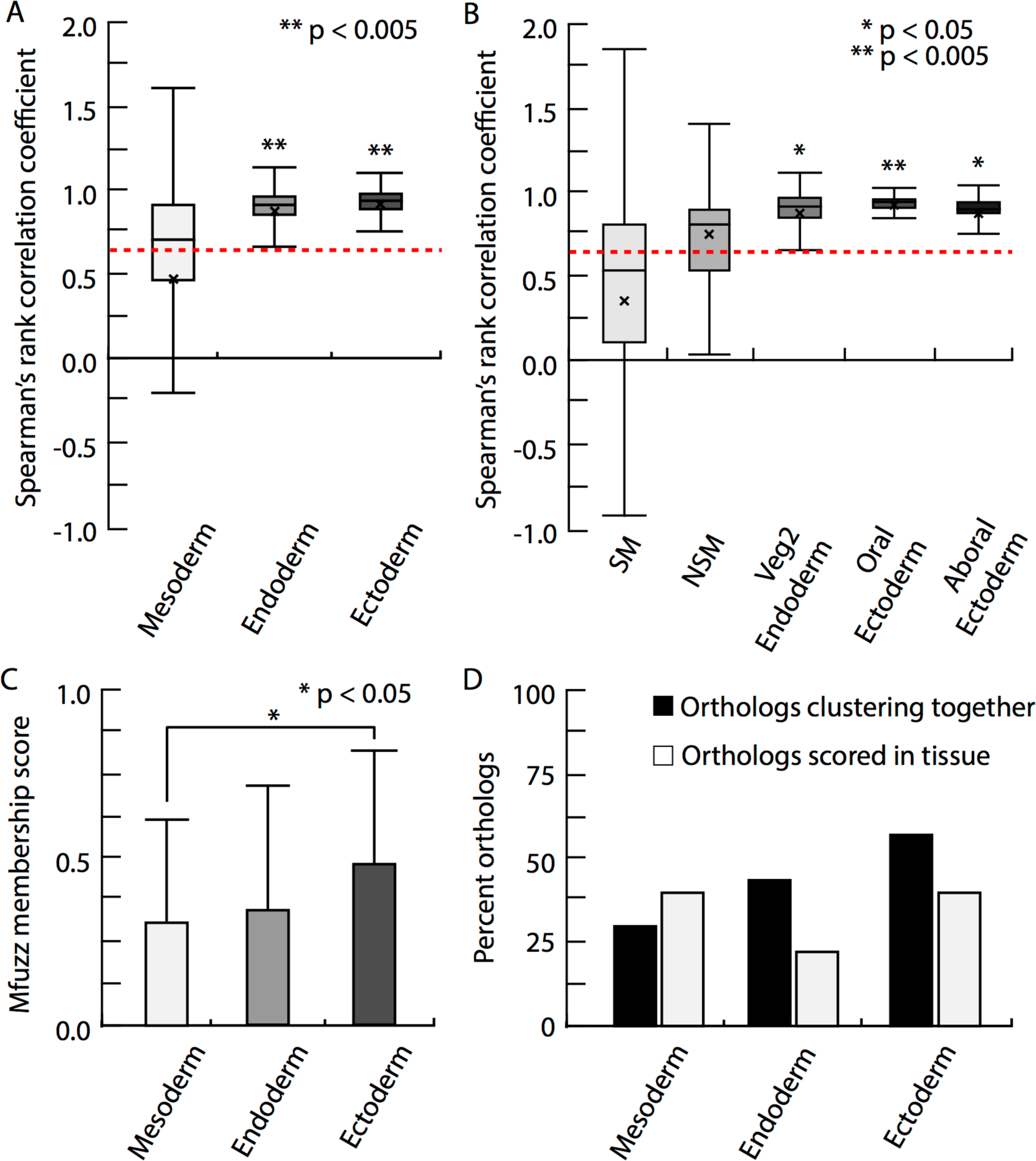
Statistical analyses of comparative timecourse data in the euechinoid *S. purpuratus* and the cidaroid *E. tribuloides.* (A-B) Distribution plots of Spearman’s rank correlation coefficients (ρ) for temporal dynamics in *E. tribuloides* and *S. purpuratus*. Genes were binned by embryonic domain of *S. purpuratus* spatial expression. Boxplot boundaries show interquartile range, means and standard deviation. Asterisks mark statistical significance as determined by a two-tailed t-test. (A) Boxplots for statistical distribution of endodermal, ectodermal and mesodermal regulatory factors in *E. tribuloides* and *S. purpuratus*. Mean ρ for endodermal and ectodermal regulatory factors were significantly higher than the mean ρ, marked as a red dashed line. Mesodermal regulatory factors did not significantly vary from the mean. (B) Boxplots for statistical distribution of subdomains of ectodermal and mesodermal regulatory factors in *E. tribuloides* and *S. purpuratus*. (C) Mean mfuzz membership scores derived from cluster analysis of regulatory orthologs. Genes were binned by their *S. purpuratus* embryonic domain of expression. Membership scores for *E. tribuloides* orthologs were binned based on their spatial distribution and compiled for statistical analysis. *E. tribuloides* ectodermal orthologs exhibited higher membership scores than mesodermal orthologs (Mann Whitney U Test, p<0.05). (D) Histogram showing both the distribution of tissue-specific orthologs examined in this study and the percent of *S. purpuratus* and *E. tribuloides* orthologs clustering together. Ectodermal and endodermal orthologs clustered together more often than mesodermal orthologs

## Discussion

### Extensive divergence of ectodermal and mesodermal regulatory linkages has occurred in echinoid developmental GRNs

This study investigated embryonic pattern formation and regulatory interactions of the euechinoid ectodermal and mesodermal GRNs in a distantly related cidaroid. By contrasting our findings with regulatory diagram of the *S. purpuratus* developmental GRN, we are able to better understand how the regulatory interactions patterning the early embryo have diverged since the cidaroids and euechinoids last shared a common ancestor in the Paleozoic 268.8 mya (Fig. 6). Numerous regulatory interactions were found to have diverged in these two clades, including (Fig 6A). Statistical analyses of comparative timecourse gene expression data revealed that gene expression profiles of regulatory gene orthologs expressed in ectodermal cell lineages are more similar to their euechinoid counterparts than those expressed in mesodermal cell lineages. This result is consistent the with the broad differences observed in cidaroids regarding SM specification, as well as that revealed here in NSM segregation and regulatory states. By contrasting these observations with those in other echinoderms, we can begin to appreciate the degree to which embryonic developmental GRNs are constrained or altered over vast evolutionary distances and can reconstruct the ancestral regulatory states that must have existed in the embryos of echinoderm ancestors (47). In the following we attempt to interpret the preceding results in an evolutionary context within the echinoderm phylum.

**Fig 6.**
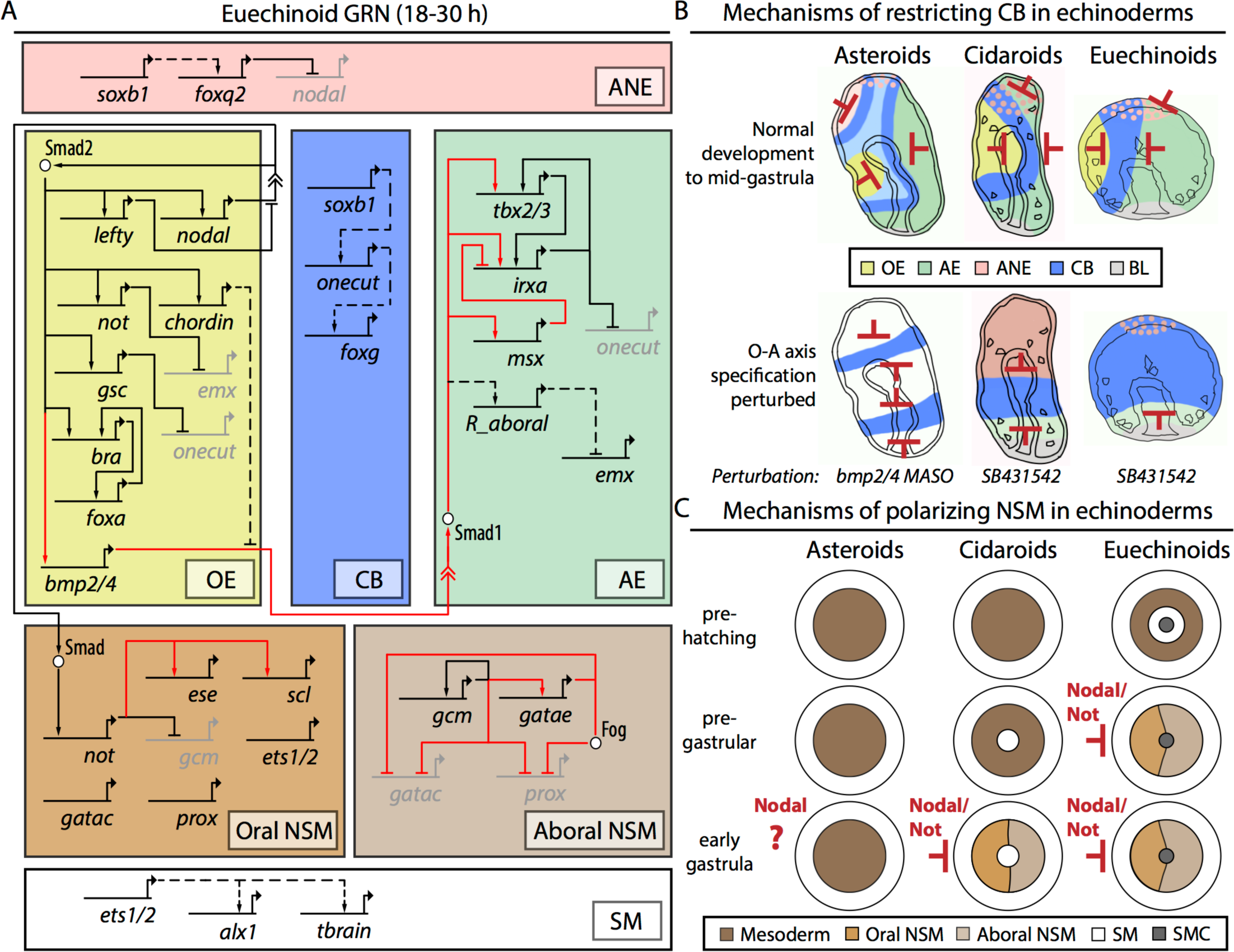
Modeling divergent and conserved regulatory transactions since the divergence of cidaroids and euechinoids. (A) A truncated *S. purpuratus* GRN showing ectodermal and mesodermal regulatory interactions investigated in this study. Solid black lines indicated conserved interactions; dashed black lines indicate interactions that are likely conserved; solid red lines indicate divergent regulatory interactions. Genes and their *cis*-regulatory regions are represented as flat lines with arrows. Embryonic domains of the echinoid embryo are represented by large boxes: pink, anterior neural ectoderm (ANE); yellow, oral ectoderm (OE); blue, ciliary band (CB); green, aboral ectoderm (AE); brown, oral non-skeletogenic mesoderm (NSM); light brown, aboral non-skeletogenic mesoderm (NSM); white, skeletogenic mesoderm (SM). (B) Regulatory states and restriction of ectodermal CB in asteroids, cidaroids and euechinoids. Red bars indicate repressor genes restrict CB in a particular embryonic domain. Color scheme is the same as in (A). The top row of schematics represents the wild-type state in three echinoderm taxa. The bottom row represents the effects of perturbation of critical ectodermal regulatory genes. (C) Mechanisms of polarization of NSM in echinoderms. Mesodermal segregation is represented in three echinoderm taxa: Asteroids, Cidaroids, and Euechinoids. In both cidaroids and euehinoids, Nodal signaling via the transcription factor Not segregates oral-aboral NSM cell types by restricting regulatory factors to the aboral side of the embryo. Whereas asteroids show no indication of NSM oral-aboral polarity, cidaroids and euechinoids lineages have undergone dramatic heterochronic shifts since the divergence of these two clades, resulting in the diversification of NSM regulatory states.

### Divergent regulatory interactions in the evolution of echinoid ectodermal and mesodermal GRNs

Our analysis utilized the regulatory transactions of the well-known *S. purpuratus* GRN as a basis for comparison between the cidaroid *E. tribuloides* and euechinoids (Fig. 6A). The comparative approach combined with the well-documented fossil record of echinoids allowed us to reveal a first approximation of the magnitude of change incurred by these GRNs since these two clades last shared a common ancestor. Spatiotemporal expression patterns of ectodermal regulatory genes in *E. tribuloides* and euechinoids strongly suggest that alteration to this circuitry is nontrivial in early development relative to the circuitry directing development of mesodermal lineages. However, while numerous regulatory transactions are conserved in the ectodermal GRN, deployment and rewiring of circuitry during the evolution of euechinoid lineages that possess direct-developing, non-feeding larvae has occurred frequently (52, 78–80). And as we have seen from this study, our data indicate that the ectodermal GRN has undergone at least one significant alteration since the cidaroid-euechinoid divergence, viz. the *bmp2/4, tbx2/3* and *irxa*. These observations support the idea that the constraints on alteration of GRN circuitry are not sufficient to maintain these linkages over vast evolutionary timescales. For instance, perturbation of Nodal signaling revealed that, while initial specification events are highly similar, alterations likely have occurred to the regulation of *bmp2/4* and *tbx2/3*. In *E. tribuloides tbx2/3* is expressed in AE and aboral NSM by mid-gastrula. By late gastrula, it is expressed in the lateral clusters of skeletogenic synthesis, at the tip of the gut, in the gut endoderm, and residually in the ectoderm. This unfolding pattern of *tbx2/3* expression in *E. tribuloides* has essentially been compressed into the early stages of euechinoid development (60). In euechinoids, perturbation of Nodal signaling with SB431542 extinguishes aboral ectodermal *tbx2/3* specifically in *P. lividus*, while not affecting its expression in SM (40). In *E. tribuloides* we observed the expression domain of *tbx2/3* expand into OE upon perturbation with this inhibitor (Fig. 2B and S5C). This observation combined with the result that *bmp2/4* responds differently to Nodal perturbation suggests altered GRN circuitry downstream of Nodal.

### Evolution of ciliary band patterning mechanisms in echinoderms

Ciliary band formation and ANE patterning in *E. tribuloides* are evolutionarily interesting given their position in the echinoderm phylogeny and given that cidaroids lack the pan-deuterostome apical senory organ (12, 13, 81, 82). Previous studies indicated that ANE patterning in *E. tribuloides* is more similar to outgroup echinoderms than it is in euechinoids; though expression of CB and anterior regulatory factors, e.g. *onecut* and *nk2.1,* exhibited spatial distributions similar to those seen in euechinoids (75). Here, we observed patterning and regulation of CB that are consistent with the hypothesis that this process is conserved in echinoids (Fig. 6B). Additionally, we observed the sequential spatial restriction of *foxq2* to ANE, a pan-bilaterian observation driven by endomesodermal *wnt* factors (23, 83–86). These data suggest that specification of the apical sensory organ in *E. tribuloides* is developmentally downstream of these events and that the loss of this embryonic structure had little effect on conserved patterning of CB and anterior localization of *foxq2*.

### Regulatory states and polarity of NSM in *E. tribuloides*

Examining mesodermal polarity in euechinoids and outgroup echinoderms aids in the establishment of a timeline of GRN evolution. In euechinoid mesodermal NSM, *gcm* is directly downstream of Notch signaling and later is restricted prior to gastrulation to aboral NSM by Not, which is directly downstream of Nodal/SMAD signaling (37, 71, 87, 88). In cidaroids, early expression of *gcm* is not dependent on Delta/Notch [Erkenbrack, Davidson and Peter, forthcoming] and its mesodermal polarity is only apparent 4-6 hours after gastrulation has begun. Thus, if *gcm* is near the top of the NSM GRN in cidaroids (16), as is the case in euechinoids (88), then this pregastrular NSM polarization can be viewed as a euechinoid synapomorphy. This hypothesis is supported by the observation that no significant polarity occurs in mesodermal specification of asteroids and holothuroids (89). Here we showed that the polarization of *gcm* and *ese* also seems to be a derived feature of echinoids and the mechanism of NSM segregation via Nodal/SMAD signaling is likely conserved (Fig. 6C). Two observations make *gcm* polarization via the transcription factor *not* (27) likely to exist in *E. tribuloides*: (1) *not* expression is observed in the oral side of the archenteron by early gastrula stage when *gcm* is spatially restricted (Fig. 1J4 and 1L3) and (2) aboral localization of *gcm* does not obtain when O-A axis patterning, and thereby *not* expression, is perturbed (Fig. 2B and S5C). Together these observations suggest a conserved role for OE regulatory factors in patterning the NSM of echinoids and that, in the lineage leading to modern euechinoids, deployment of GRN circuitry polarizing NSM underwent a heterochronic shift in the lineage leading to euechinoids.

The data presented on O-A polarity in the NSM of *E. tribuloides* suggest that multiple regulatory domains unfold at and around the tip of the archenteron as gastrulation proceeds. This is also the case in euechinoids (90). However, this study determined that *ese* operates in the oral NSM exclusive of *gcm* in the aboral NSM. While the regulatory states in *E. tribuloides* NSM need further refinement by two-color WMISH, for our purposes the overt disorder in its formation relative to the overt order of *S. purpuratus* NSM makes two salient points. First, early pregastrular or early gastrular polarity of NSM regulatory states represents an echinoid novelty, as no evidence for early mesodermal polarity exists in outgroup echinoderms (89, 91). Second, if we take *E. tribuloides* as a proxy to the ancestral state for this character/regulatory state, then it is clear that the O-A polarity observed in the euechinoid NSM was shifted to occur prior to gastrulation in the lineage leading to modern euechinoids. On the other hand, an alternative evolutionary scenario is that NSM polarity manifested in these two modern echinoids is the result of two independent evolutionary trajectories with heterochronic and spatial differences, but both meeting a similar end in the diversification of NSM cell types in early development. That at least two O-A regulatory states are common to these embryos and that *gcm* is downstream of Nodal and much later is downstream of Notch signaling provide support for the first scenario. Further investigation into the developmental timing and regulatory states of NSM segregation in *E. tribuloides* and other cidaroids is required to parse out the most likely evolutionary scenario of NSM evolution in echinoids.

### Evolution of global embryonic domains in early development of echinoids

Previous analyses of embryonic regulatory states in *E. tribuloides* surveyed SM regulatory factors (17, 47) and anterior neural ectoderm specification (75). Additionally, two previous studies investigated SM and early endomesodermal micromere regulatory factors in the Pacific-dwelling cidaroid *Prionocidaris baculosa* (16, 92). Integrating these data into this study affords an analysis of global embryonic regulatory states and GRN linkages over 268.8 mya of evolution in indirect-developing sea urchins. From these studies, numerous alterations to deployment and GRN circuitry at all levels of GRN topology can be enumerated. Here, we enumerate 19 changes in spatiotemporal deployment or regulation of ectodermal and mesodermal embryonic regulatory factors since the cidaroid-euechinoid divergence (Table S1). Prominent among alterations of regulatory interactions are those that have occurred in establishing polarity in mesodermal embryonic domains. Endodermal and ectodermal specification and regulatory states also have undergone significant change, but to a lesser degree. One hypothesis that can accommodate these observations is that endodermal and ectodermal developmental programs may be more recalcitrant to change than mesodermal programs due to their more ancient evolutionary origin (5), suggesting that accretion of process over evolutionary time is a mechanism of constraint in developmental programs (5). Indeed, in euechinoids there have been additional layers of GRN topology accrued in mesodermal specification, e.g. the *pmar1-hesc* double-negative gate novelty (16, 17, 22), delta-dependent NSM specification (17, 34), etc., which cidaroids do not exhibit, and which may explain the observation that little to no appreciable change has been observed in the mesodermal developmental programs of *L. variegatus, P. lividus* and *S. purpuratus,* representatives of modern euechinoid lineages that diverged approximately 90 mya.

Since the divergence of cidaroids and euechinoids, numerous alterations to developmental GRNs have accrued in these lineages. This study revealed that changes to mesodermal GRN architecture have occurred more frequently than alterations to ectoderm GRN architecture. These results support the notion that GRN architecture evolves at different rates (93) and provide an in principle explanation for the rapid evolution observed in both cidaroid and euechinoid sea urchin lineages that have independently evolved direct-developing larval forms (53). It remains to be determined by future research efforts whether the shared regulatory states between cidaroids and euechinoids elucidated here are the product of conserved stretches of genomic DNA hardwired in the *cis*-regulatory regions of the orthologous regulatory genes or whether they are the result of diverged *cis-* regulatory modules producing similar developmental outcomes.

### Materials and Methods

Animal and embryo culture, cloning, acquisition of spatial and temporal gene expression data and microinjection perturbation data were as previously described (17, 47). Dosage of small molecule inhibitors DAPT and SB431542 were determined by dilution series. Concentrations for DAPT and SB431542 were 12 µM and 15 µM respectively. Additional details for previously described methods, statistical analyses and all other experimental manipulations can be found in SI Materials and Methods.

## Acknowledgments

We appreciatively acknowledge numerous colleagues for discussing various aspects of this work with us over the long course of its maturation: Dr. Julius C. Barsi kindly provided his knowledge and the *Sp onecut* engineered BACs; Jennifer Wellman, M.S., for her insight on statistical analyses; Stefan Materna provided raw data for *S. purpuratus* mRNA transcript abundance and reviewed early versions of the manuscript; Matthias Futschik provided mfuzz analysis advice; Drs. Oliver Griffith, David McClay, Tom Stewart and Jeffrey R. Thompson provided critical commentary on figures and the manuscript. The commentary of two anonymous reviewers greatly improved data analysis and the manuscript. This work was developed and written in the laboratory of Professor Günter P. Wagner; his patience and support are fondly appreciated. This work was funded by National Science Foundation CREATIV grant #1240626.

